# AlphaFold3 for Non-canonical Cyclic Peptide Modeling: Hierarchical Benchmarking Reveals Accuracy and Practical Guidelines

**DOI:** 10.1101/2025.05.24.655528

**Authors:** Chengyun Zhang, Wentong Wang, Ning Zhu, Zhigang Cao, Qingyi Mao, Cheng Zhu, Chenhao Zhang, Jingjing Guo, Hongliang Duan

## Abstract

Despite the revolutionary impact of AlphaFold3 on structural biology, this model’s capability in predicting non-canonical cyclic peptides remains unexplored. Given the clinical significance of cyclic peptides containing unnatural residues as a therapeutic modality, we present the first systematic evaluation of AlphaFold3 for this class of molecules. To facilitate benchmarking, we developed an automated input pipeline to streamline AlphaFold3 predictions for cyclic peptides. Our study aims to: (1) quantify the hierarchical accuracy (all-atoms, Cα atoms, and atoms of unnatural residue levels) of AlphaFold3 in predicting both non-canonical cyclic peptide monomers and complexes; (2) assess the reliability of AlphaFold3’s confidence metrics; (3) evaluate the influence of multiple sequence alignment (MSA) and structural templates; and (4) identify systematic biases in AlphaFold3’s predictions. Based on these analyses, we provide practical guidelines for applying AlphaFold3 in cyclic peptide structure prediction for facilitating the related research of bioactive cyclic peptides.

## Introduction

The accurate prediction of biomolecular structures has been transformed by deep learning approaches, particularly since DeepMind’s introduction of AlphaFold in 2019, a groundbreaking framework for the long-standing challenge of protein structure determination^1^. This innovative approach leverages multiple sequence alignments (MSA) and templates and harnesses the power of attention mechanisms to enable fully automated and highly accurate protein structure prediction. This contribution was recognized by the 2024 Nobel Prize in Chemistry. The original AlphaFold was designed for monomeric protein structure prediction. Inspired by its remarkable success, the scientific community sought to adapt this model for structure prediction including multiple chains, employing strategies such as linker incorporation or sequence extension. Responding to growing scientific demand, DeepMind subsequently released AlphaFold-Multimer^2^ in 2021, which expanded capabilities to protein complexes including multimeric protein assemblies and nucleic acid interactions. In addition, the AlphaFold-based advancements spurred the development of complementary prediction systems^3–8^ like RoseTTAlphaFoldold^3^. Moreover, it also revolutionized protein design through tools^9–15^ such as AlphaFoldDesign^10^ and BindCraft^15^, enabling reasonable generation of novel structural motifs with great potential.

Despite these transformative developments, current implementations of both AlphaFold and AlphaFold-Multimer remain limited in their ability to accurately predict the structures of cyclic peptides, an important class of bioactive molecules with unique cyclic structural constraints and therapeutic potential^16^. The clinical significance of this constrained cycle feature is exemplified by oxytocin, a cyclic peptide administered to approximately 50% of the over 3 million annual childbirth cases in the US alone^17^. However, neither AlphaFold nor AlphaFold-Multimer can reliably predict their structures, particularly for variants containing non-canonical amino acids.

The first attempts to address this gap is proposed by the Bhardwaj group who enabled AlphaFold model to predict cyclic peptides containing natural amino acids through the incorporation of cyclic constraints into its positional matrices^13^. Based on their work, our team subsequently made the optimizations to improved prediction accuracy for natural cyclic peptides^18^. However, the fundamental limitations remained for non-canonical cyclic peptide variants. Although traditional computational approaches^19–21^ like Rosetta^19^ can partially address this problem, their prohibitive computational resources, unsatisfactory accuracy and extensive expert tuning limit their practical utility.

The recent release of AlphaFold3 represents a potential breakthrough through its novel residue and atom-level combined feature engineering, enabling the prediction of modified peptides and small molecules^22^. Notably, its architecture also supports the cyclic peptide prediction. However, the majority of benchmarking^23–25^ have primarily evaluated its predictive accuracy for linear structures. Its performance on non-canonical cyclic peptides remains unexplored, posing risks of experimental bias if applied without validation.

Given the unexplored potential of AlphaFold3 for non-canonical cyclic peptides, this study aims to present the first comprehensive evaluation of this model for non-canonical cyclic peptide prediction. To facilitate high-throughput analysis, we developed an automated pipeline for AlphaFold3 input preparation, eliminating the time-consuming manual data entry required by the graphical web interface. Through systematic benchmarking of diverse structures, we:(1) quantify prediction accuracy across multi-layered structural features, including all atoms, Cα atoms, and atoms of unnatural residues (2) identify systematic biases of AlphaFold3. (3) investigate the impact of MSA and templates on model performance. (4) assess the reliability of confidence metrics. Furthermore, our findings provide essential guidelines for applying AlphaFold3 to non-canonical cyclic peptide structure prediction and reveal fundamental insights into both its capabilities and limitations.

### Data

The cyclic peptides including unnatural amino acids used in this study, were sourced from three data sources: (1) the cPEPmatch webserver^26^, (2) the Protein Data Bank (PDB)^27,28^, and (3) a designed structure set from Baker’s research^29^ focusing on membrane-traversing macrocycles. Initial data selection includes retaining only structures with head-to-tail cyclization or disulfide bridge formation and excluding those that have cyclic peptide residues exceeding 50. Following this collection, all structures underwent curated validation using PyMOL (version 3.1.3.1, Schrödinger LLC) to verify their integrity. Each entry was manually annotated with its release year, residue length, and secondary structure features et. al information. Finally, this curation process yielded 85 high-quality structures, including 68 monomer and 17 complex forms.

The monomer subset contains samples with residue lengths ranging from 5 to 43, including 4 α-helix, 21 β-sheet, 40 loop, and 3 mixed-type secondary structures. As for the complex subset, it comprises samples with ligand residue lengths from 6 to 15, featuring 5 α-helix, 10 β-sheet and 2 loop conformations. Figure 1B illustrates the structural diversity and cyclic peptide residue length distribution of our dataset. Complete dataset specifications, including PDB accession codes and related information are available after publication.

**Figure 1.**
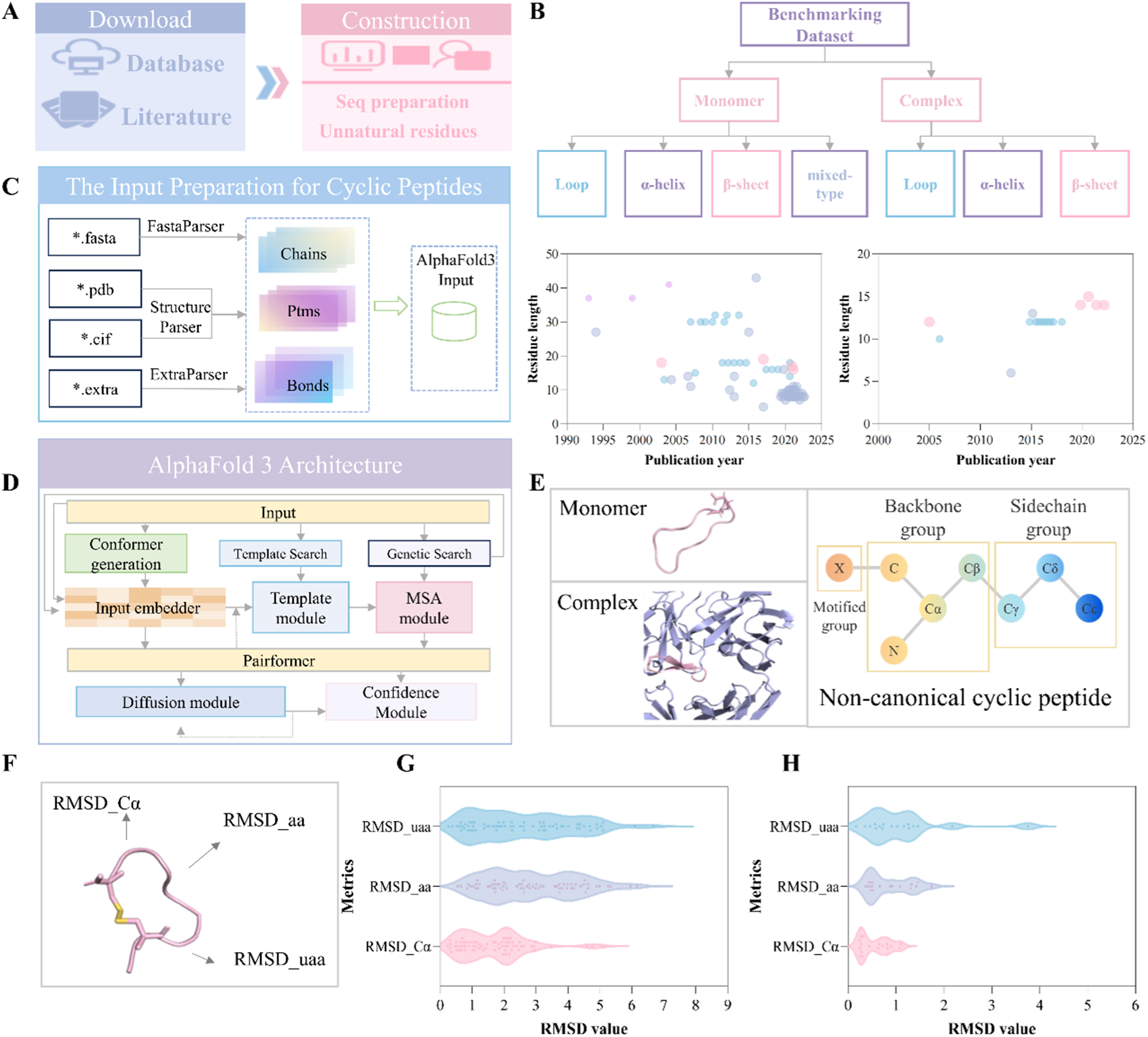
Overview of the benchmarking study. (A) Construction of the benchmarking dataset: data download and filtering. (B) Detailed information about the benchmarking dataset: secondary structure types, residue length, and publication year. (C) AlphaFold3 input preparation for cyclic peptide structure prediction. (D) Architecture of the AlphaFold3 model. (E) AlphaFold3 predicted structure for cyclic peptides: atom-level and residue-level features. (F) Hierarchical RMSD evaluation for cyclic peptide structures. (G) Value distribution of RMSD_Cα, RMSD_aa, and RMSD_uaa for predicted cyclic peptide monomers. (H) Value distribution of RMSD_Cα, RMSD_aa, and RMSD_uaa for predicted cyclic peptide ligands in complexes.

## Method

### Model

The benchmarking experiments were performed on a workstation running Ubuntu 20.04.2. The system configuration comprised an Intel Xeon Gold 6430 CPU (64 cores), four NVIDIA A800 GPUs (80 GB memory each; 320 GB total GPU memory), 500 GB of RAM, and 40 TB of storage space. For structure prediction of cyclic peptides, we utilized the open-source AlphaFold 3 implementation (available at: https://github.com/google-deepmind/alphafold).

### Metric

In the benchmarking experiments, we evaluated the performance of the AlphaFold3 model in predicting non-canonical cyclic peptide monomer and complex structures. Two scoring systems were implemented to comprehensively assess the predicted structures.

For the monomer prediction task, we employed three RMSD-based metrics to evaluate different aspects of the predicted cyclic peptide structures:

- RMSD_aa: Computes the deviation of all atoms in the cyclic peptides.
- RMSD_Cα: Measures the deviation of Cα atoms in the cyclic peptide backbone chains.
- RMSD_uaa: Specifically assesses the deviation of atoms of unnatural amino acids.

These metrics collectively provided a comprehensive evaluation of AlphaFold3’s performance in modeling the overall structure, backbone chains, and unnatural residue components of cyclic peptide monomers, thereby exploring the performance of the AlphaFold3 model in predicting cyclic conformation. The calculation for RMSD metrics is detailed below:

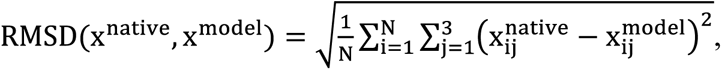

where x^native^ is the native cyclic peptide structure, x^model^ is the predicted cyclic structure provided by the AlphaFold3, index i is the number of atoms (all atoms, Cα atoms, atoms of unnatural amino acid) N, and index j is the 3D coordinates.

In addition to RMSD-based metrics, we computed confidence scores including the predicted local-distance difference test (pLDDT) and predicted TM-score (pTM) to evaluate model reliability. These metrics were employed to analyze their correlation with structural deviations and assess their effectiveness in validating the accuracy of predicted conformations. Among these, pLDDT, a widely adopted indicator, was used to reliably predict the local-distance difference test accuracy of the corresponding structure. The LDDT is a superposition-free metric that assesses the local structural accuracy of atomic models by comparing interatomic distance differences, while simultaneously validating stereochemical plausibility. AlphaFold3 provide a version of LDDT that considering distances between all atoms and polymer residues peptide residues. The LDDT score for atom *l* computed as:

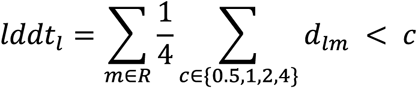

Where d_lm_ denotes distance between atom *l* and atom m in the AlphaFold3’s mini-rollout prediction, *l* denotes all atoms and the set of atoms *m* ∈ R that satisfying (1) the distance in the native structure between atoms *l* and m < 15 Å (if *m* is a protein atom) or < 30 Å (if m is a nucleic acid atom). (2) exclusively polymer chain atoms;. (3) One atom per token (Cα for standard protein residues and C1′ for standard nucleic acid residues).

As for the pTM, it is a confidence measure based on the predicted histogram and orientations to predict template modeling score^30^ that indicates the global quality of structure. The calculation of TM-score is shown below:

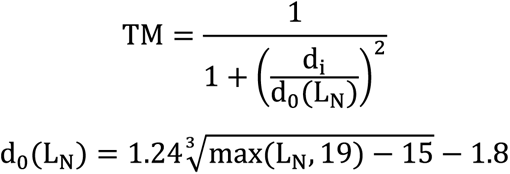

where L_N_ denotes the number of amino acids in the native structure, d_i_ is the Euclidean distance between the aligned amino acid in the predicted structure and its corresponding residue in the native structure, and d_0_(L_N_) is a distance scale factor, empirically determined based on expert knowledge as described in the paper.

For the complex prediction task, we evaluated the cyclic peptide ligand with RMSD_aa, RMSD_Cα, and RMSD_uaa, as we did in the non-canonical cyclic peptide monomer task. Although the RMSD-based metrics effectively quantify the structural similarity between the predicted and native structures for both the ligand and the target, they do not explicitly indicate whether the predicted ligand is correctly positioned at the binding site of the protein. Therefore, the fraction of native contacts (F_nat_)^31^ metric was used to address this limitation by measuring the fraction of native interfacial contacts preserved in the predicted complex interface. The reliability of complexes is classified according to F_nat_ using the following criteria: (1) high-accuracy predictions (F_nat_ > 0.8), (2) medium-accuracy predictions (0.5 < F_nat_ ≤ 0.8), (3) acceptable predictions (0.2 < F_nat_ ≤ 0.5), and (4) incorrect predictions (F_nat_ ≤ 0.2). The F_nat_ metric is calculated as follows:

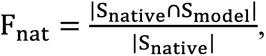

and

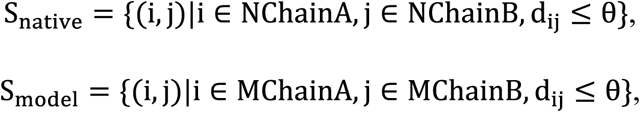

where S_native_ represents the set of atom pairs (one from the protein target and one from the ligand) in the native structure with a distance less than threshold θ, S_model_ represents the set of atom pairs (one from the protein target and one from the ligand) in the predicted structure with a distance less than threshold θ, operator ∩ is the intersection of two sets, operator |.| indicators the size of a set. Two atoms on different sides of the interface can be regarded to be in contact if their distance is less than 5 Å.

Multiple confidence metrics were employed to assess the reliability of predicted complexes. In this study, we systematically evaluated the correlation between these confidence scores and structural accuracy (as measured by RMSD) to determine their predictive value. Building upon our methodology for cyclic peptide monomer evaluation, we specifically examined the ligand_pLDDT metric for complex prediction assessment. Additionally, instead of focusing on entire protein chains, we primarily employed interface-specific metrics. One key metric is ipTM (interface predicted TM-score), which evaluates the structural similarity of predicted interfaces to the native structures. The calculation of ipTM follows the same methodology as described above we mentioned. Additionally, we computed the ipAE (interface predicted aligned error) for cyclic peptide complexes in our benchmarking experiments.

pAE^2^ is a measure of the confidence of the relative position between two residues within the predicted structure, where aligned error e_ij_ is defined as the error at residue j if the predicted and true structures are aligned using the backbone frame of residue i, formulated as follows.

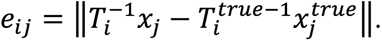

where *x_j_* and 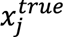 are the predicted and true local coordinates of residue j, 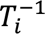 and 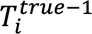 are the optimal transformation inverse matrix of coordinates for the predicted and true structures when aligning based on residue i. During model training, the aligned error can be calculated according to the above formulation given the predicted and true structures. Then the distribution of aligned error is discretized into 64 bins, covering the range from 0 to 32 Å with 0.5 Å bin width, to represent it using a one-hot vector. At the same time, the probability distribution of aligned error can be readily predicted by a linear projection of the residue embedding, followed by a softmax operation. Thus, the aligned error is optimized by the categorical cross-entropy loss between the predicted and true distributions. This metric quantifies the confidence in the relative positioning of residues within the predicted interface, with lower values indicating higher reliability.

## Results

Despite AlphaFold3 has demonstrated remarkable capabilities in the non-canonical peptide/protein structure prediction, the lack of systematic benchmarking and conclusive evidence regarding its feasibility and accuracy for predicting non-canonical cyclic peptides and their complexes has left potential users uncertain. To address this gap, we constructed a comprehensive benchmarking dataset for non-canonical cyclic peptides and employed multiple evaluation metrics to assess the performance of AlphaFold3 in this specific task. Our study provides in-depth insights into the model’s efficacy, preferences, and limitations when applied to non-canonical cyclic peptides, offering a critical foundation for future research and practical implementation.

### AlphaFold3 Input Preparation for Cyclic Peptide Prediction

Although the AlphaFold server offers an intuitive graphical interface, its functionality is limited for large-scale cyclic peptide predictions. Researchers must spend considerable time manually inputting necessary information, highlighting the need for an automated input-preparation pipeline. To address this, we developed a streamlined preprocessing pipeline for our benchmarking dataset, enabling efficient structure prediction for cyclic peptides. This tool is also adaptable for broader cyclic peptide prediction tasks.

The tool supports multiple input file formats, including .pdb, .cif, and .fasta files. The initial processing step involves extracting chain information, non-canonical amino acid modifications, and bond information from these input files. The detailed workflow is presented in Algorithm 1. For structural files (.pdb and .cif formats), the ParseStructureFile module utilizes Biopython to parse the structure and retrieve chain information along with amino acid modification data. In the case of sequence files (primarily .fasta format), chain information and modification details are extracted through string parsing and regular expression matching. Unnatural peptide sequences should be formatted as “XXXX(XXX)XXX”, where the parenthetical component contains the Chemical Component Dictionary (CCD) code^32^ of the unnatural amino acid.

Since .pdb, .cif, and .fasta files typically lack bond information, we have implemented a supplementary file system with the .extra extension to specify both bond connectivity and additional modification details. Our standardized bond notation follows the format: [chain_id][residue_index][atom]-[chain_id][residue_index][atom]. This input mode enables accurate representation of different covalent bonds.

#### Algorithm 1 InputParser

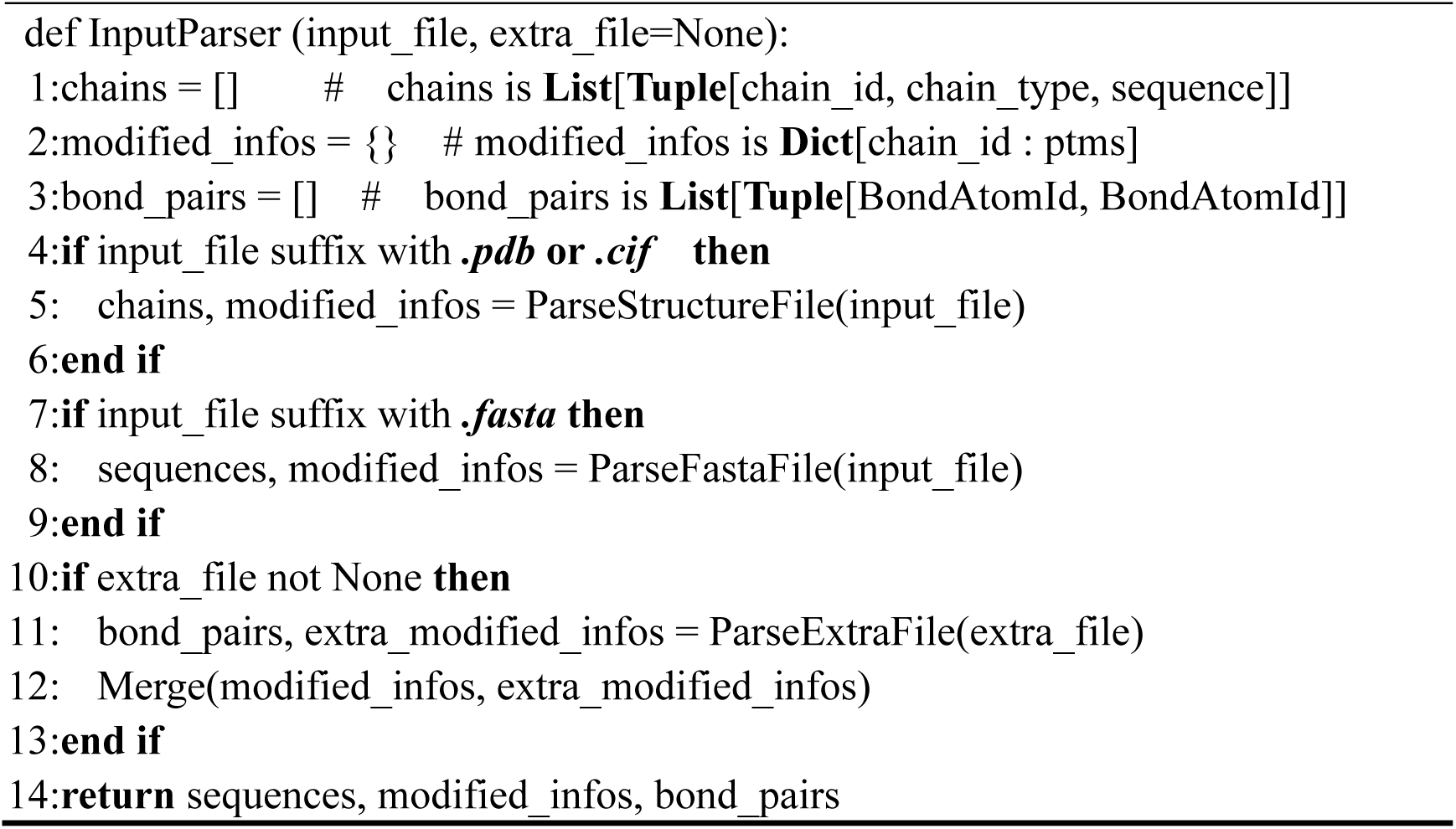

#### Algorithm 2 MockAlphaFold3Input

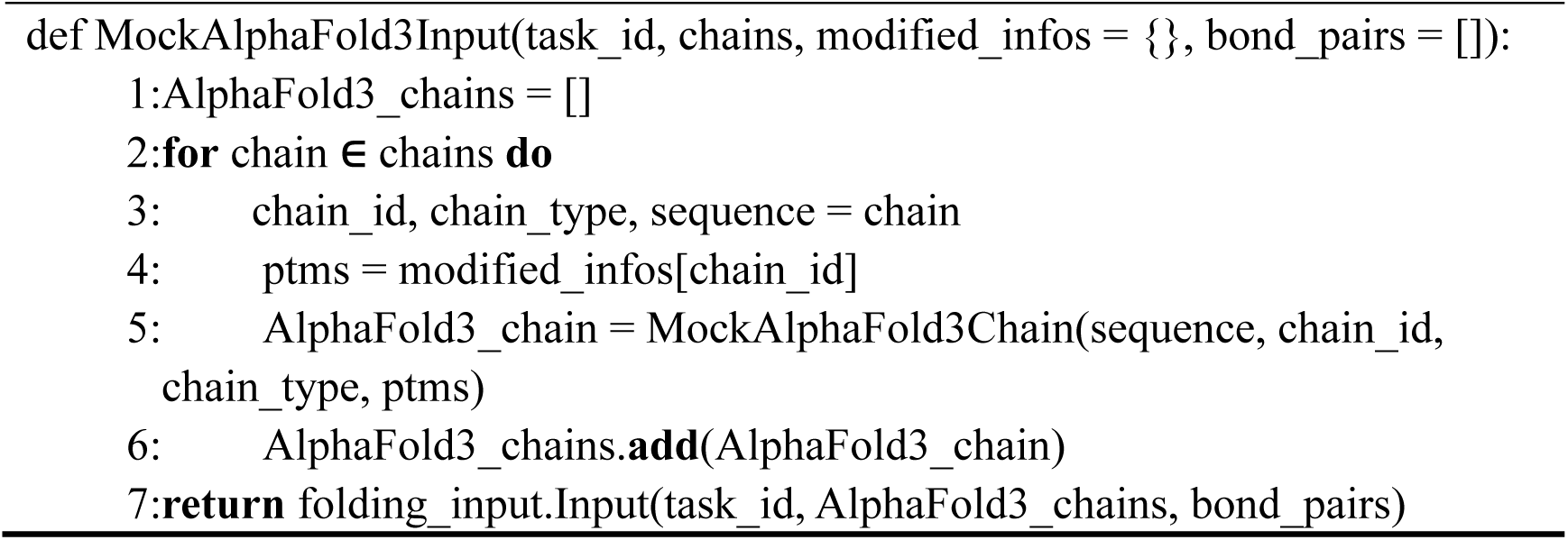

The second step involves constructing input objects compatible with AlphaFold3 based on the extracted information from the input files. As outlined in Algorithm 2, for each chain, we retrieved its chain ID and indexed the corresponding modification details from the amino acid modification dictionary. Depending on the enumerated chain type, we instantiated the appropriate chain object as defined by AlphaFold3. Finally, we generated a usable input object by initializing folding_input.Input from AlphaFold3. For standard AlphaFold3 JSON input files, the AlphaFold3 source code’s pipeline includes a built-in load method that directly converts the JSON data into a folding_input. Input object. Since our pipeline simply invokes this existing procedure, we will not elaborate further on the processing of .json files for the sake of brevity.

After completing the input preparation, we conducted the benchmarking experiment based on monomer and complex datasets to systematically evaluate the accuracy and potential biases of AlphaFold3 in predicting cyclic peptide structures. The detailed analyses are shown below.

### The Evaluation of AlphaFold3-Predicted non-canonical Cyclic Peptide Monomers

#### The RMSD-based evaluation for non-canonical cyclic peptide monomers

First, we evaluated AlphaFold3’s ability to model non-canonical cyclic peptide monomers using three RMSD-based metrics. As shown in Figure 1G, the RMSD_Cα values ranged from 0.231 to 5.072 Å, indicating that the model could effectively predict cyclic peptide backbones. A similar trend was observed in the RMSD_aa values (0.552-6.164 Å).

As we mentioned before, a key strength of AlphaFold3 lies in its ability to model unnatural or modified residues. To evaluate this, we calculated the RMSD_uaa value of the predicted structure, which focuses exclusively on deviations in these unnatural residues. The RMSD_uaa values ranged from 0.261 to 6.670 Å, highlighting AlphaFold3’s accuracy in predicting non-canonical structural features. In addition, the average RMSD_Cα, RMSD_aa and RMSD_uaa achieved 1.671, 2.862 and 2.592 Å, respectively (Figure 2A), suggesting that AlphaFold3 performs better in predicting stable backbones than flexible side chains.

**Figure 2.**
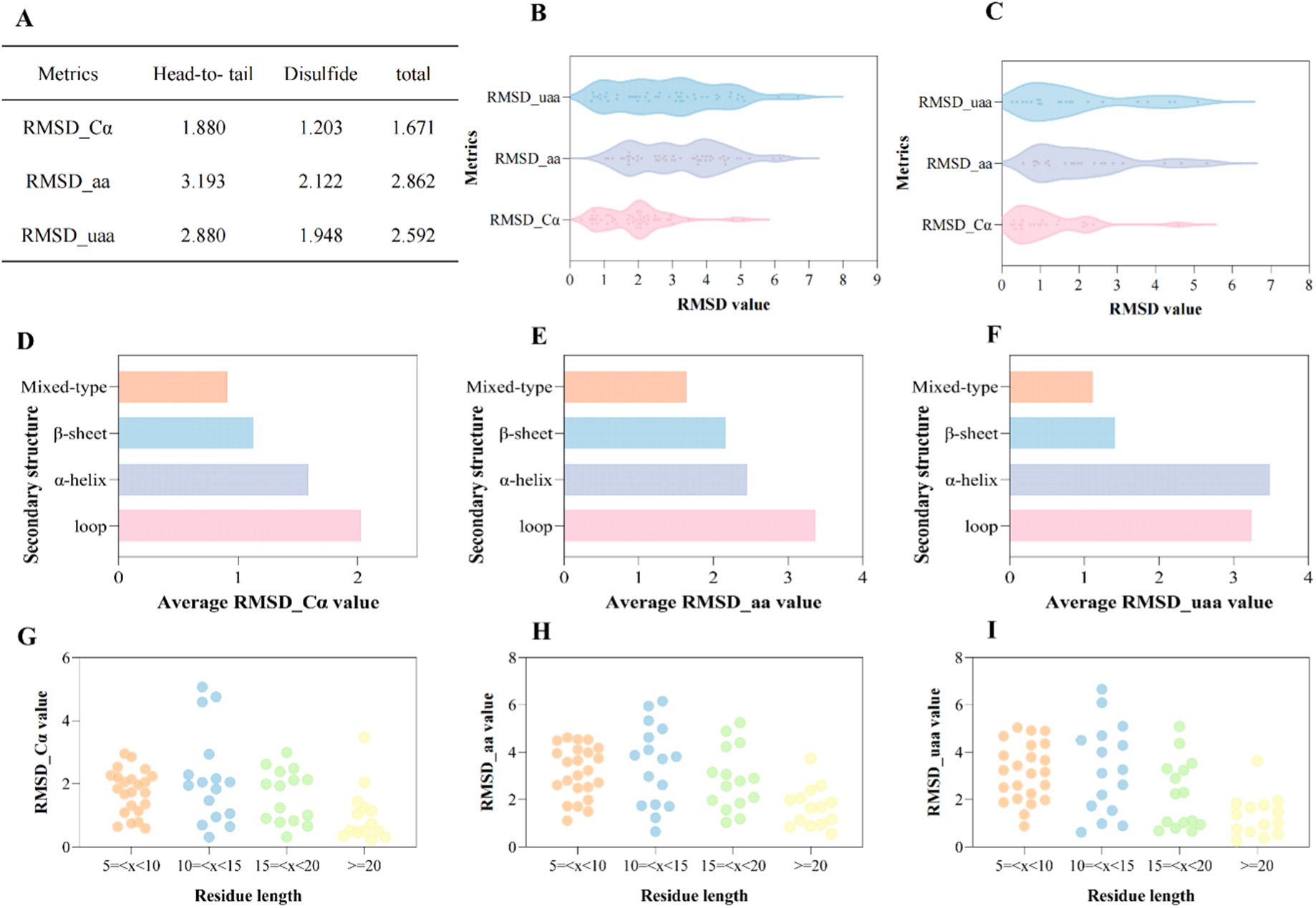
Performance of AlphaFold3 in predicting non-canonical cyclic peptide structures. (A) Average RMSD values (RMSD_Cα, RMSD_aa, and RMSD_uaa) for head-to-tail and disulfide cyclic peptide monomers. (B) Distribution of RMSD values for head-to-tail cyclic peptide monomers. (C) Distribution of RMSD values for disulfide cyclic peptide monomers. (D-F) Correlation between secondary structural elements (α-helix, β-sheet, mixed-type, and loops) and RMSD metrics. (G-I) Relationship between residue length and RMSD metrics.

When considering all atoms of the structure, subtle changes in the atoms of unnatural residue are difficult for AlphaFold3 to capture compared to the stable backbone atoms, resulting in an RMSD increase of 0.921 Å compared to the Cα atoms. In other words, although the AlphaFold3 supports the unnatural residue modeling, its performance is not as good as that of natural residues. People should carefully estimate the plausibility of unnatural residues of predicted structures.

Second, we observed a structure preference of AlphaFold3 in the disulfide cyclic peptides. As shown in the Figures 2B and 2C, the average RMSD_Cα, RMSD_aa and RMSD_uaa of disulfide cyclic peptides were 1.203, 2.122, and 1.948 Å, which were lower than those of the head-to-tail cyclic peptides (1.880, 3.193, and 2.880 Å). This difference may stem from the stabilizing effect of disulfide bonds, which restrict conformational flexibility and guide the model toward more native-like folds.

Then, we investigated the relationship between these metrics and secondary structural features, as illustrated in Figures 2D-F. Compared to disordered loop structures, cyclic peptides containing secondary structural elements (α-helix, β-sheets and mixed-type structures) demonstrated significantly lower RMSD_Cα values (1.585, 1.126, and 0.908 Å, respectively) compared to the average of 2.023 Å for loop structures. This observation suggests that well-defined secondary structures may contribute to enhanced structural predictability. In addition, further analysis of the relationship between residue length and RMSD values (Figures 2G-I) reveals that longer peptide chains (>10 residues) appear to be the formation of stable secondary motifs, thereby enabling AlphaFold3 to more accurately predict native-like cyclic peptide conformations. Interestingly, this trend reversed for shorter peptides (<10 residues), which exhibit a lower average RMSD_Cα (1.756 Å) compared to intermediate-length peptides (10-15 residues). We hypothesize that the constrained conformational space in shorter peptides limits structural flexibility, reducing the complexity of prediction.

Furthermore, we identified that some predicted head-to-tail cyclic peptide structures failed to form proper cyclic conformations, instead adopting linear-like configurations. We will analyze this phenomenon in detail in the following mechanism discussion section.

### The exploration of the reliability confidence scores

In this study, we assessed the reliability of pLDDT and pTM as accuracy indicators for predicted cyclic peptide structures. As these metrics reflect overall prediction confidence, we specifically examined their correlation with RMSD_aa and RMSD_Cα values, which serve as direct quantitative measures of structural deviation from global native conformations.

The pLDDT serves as a confidence score for structural predictions. As shown in Figures 3A-B, we analyzed the relationship between pLDDT values and RMSD-based accuracy metrics (RMSD_aa, RMSD_Cα). While no significant linear correlation was observed between pLDDT scores and these structural deviation measures, the overall prediction accuracy remained high (RMSD_Cα and RMSD_aa values both below 2 Å.), with most models exhibiting pLDDT values >90. This metric can, to some extent, indicate the authenticity of the predicted conformation. When this metric is higher, the occurrence of low RMSD structures increases. Namely, the likelihood of predicted structure assembling to the native structure increases when the pLDDT values increase. For instance, the predicted structure for 2M2S (pLDDT = 95.650) is close to the native conformation, with an RMSD_Cα of 1.236 Å and RMSD_aa of 2.089 Å, underscoring the utility of high pLDDT scores as an indicator of prediction reliability despite the absence of a clear correlation.

**Figure 3.**
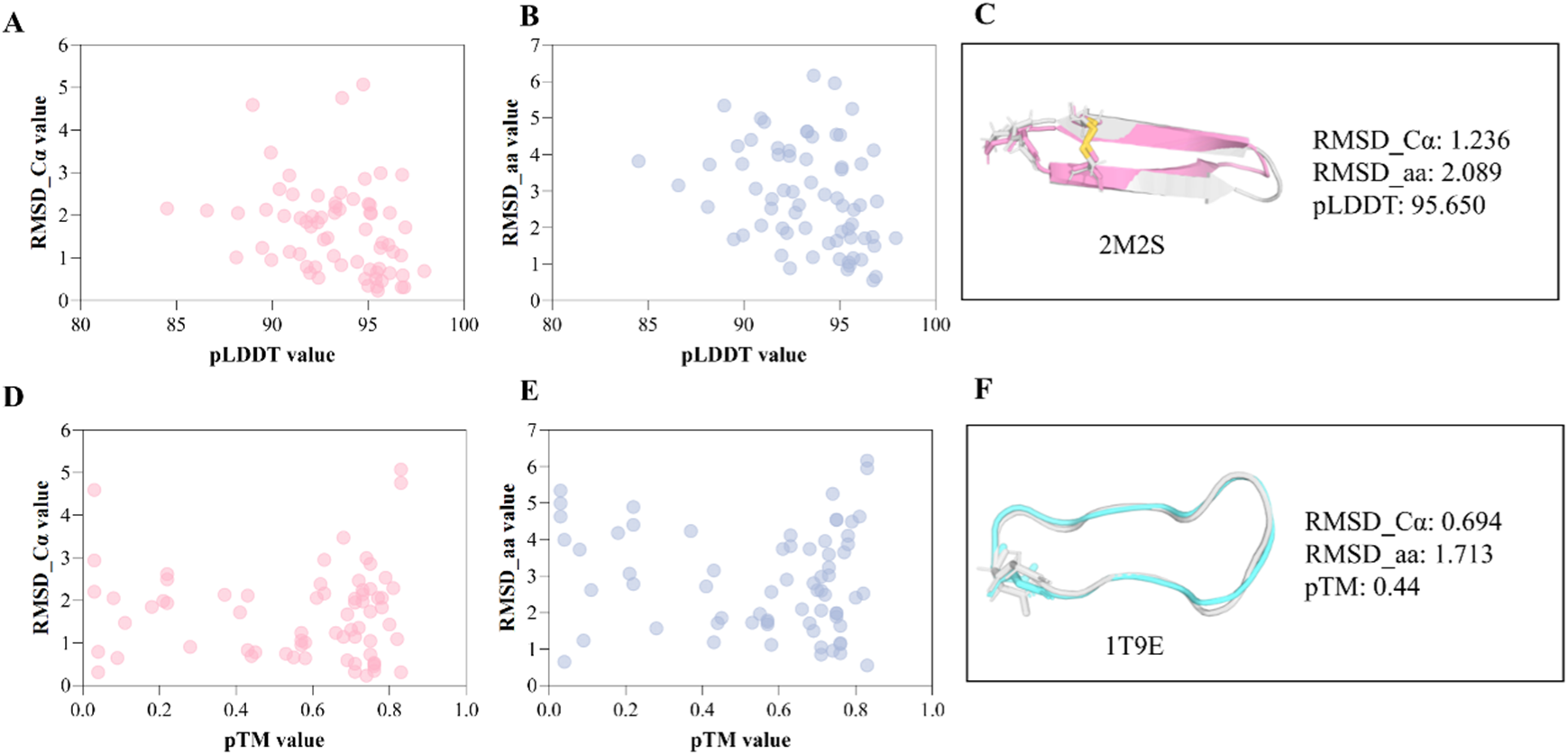
Evaluation of confidence metrics for non-canonical cyclic peptide structure predictions. (A) Correlation between RMSD_Cα and pLDDT values. (B) Correlation between RMSD_aa and pLDDT values. (C) Predicted structure of 2M2S with associated quality metrics. (D) Correlation between RMSD_Cα and pTM scores. (E) Correlation between RMSD_aa and pTM scores. (F) Predicted structure of 1T9E with associated quality metrics.

The pTM is also a popular metric for representing the confidence of this model’s generated structures, including cyclic peptide predictions. To evaluate its reliability, we analyzed the relationship between pTM values and RMSD measurements. As shown in Figures 3D-E, pTM exhibited limited correlation with the structural plausibility of predictions. Notably, even at low pTM values, the predicted structures can still achieve low RMSD_Cα and RMSD_aa values, indicating high accuracy. In the case of 1T9E (Figure 3F), the pTM was only 0.44, but the RMSD_Cα and RMSD_aa can reach 0.694 and 1.713 Å, respectively. Consequently, we recommend that users adopt pLDDT rather than pTM when evaluating the reliability of predicted cyclic peptide structures.

### The Evaluation of AlphaFold3-Predicted non-canonical Cyclic Peptide Complexes

#### The F_nat_ evaluation of non-canonical cyclic peptide complexes

As we mentioned before, the RMSD-based metrics focus more on the chain plausibility rather than the binding site accuracy. The F_nat_ is more concentrated on the interface of the contact pair, therefore implying the binding site difference between predicted and native structures. Figure 4A displays the F_nat_ values of the cyclic peptide complexes. In most cases, AlphaFold3 gave satisfactory results. Almost 70% of predicted structures were equipped with F_nat_ >0.8 and the remaining predicted structures had F_nat_ values that were lower than 0.8 Å but higher than 0.5 Å. Such results demonstrate that AlphaFold3 can effectively dock the predicted ligands and protein targets, and produce a plausible complex with high probability.

**Figure 4.**
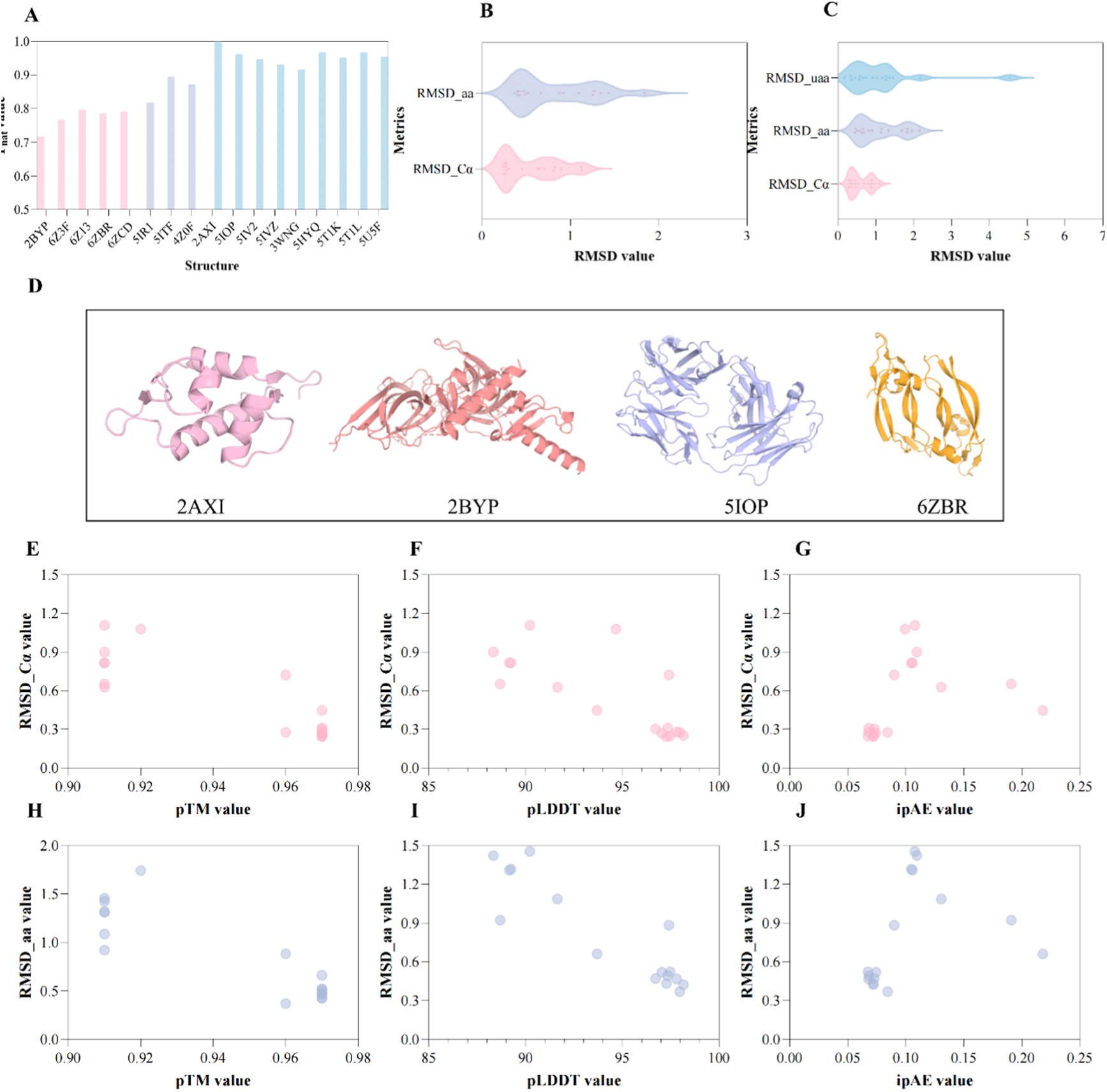
Performance of AlphaFold3 in predicting non-canonical cyclic peptide complexes. (A) F_nat_ values of predicted structure (B) Distribution of RMSD values for predicted target structure of cyclic peptide complexes. (C) Distribution of RMSD values for predicted ligand structure of cyclic peptide complexes. (D)The different target classes. (E-G) Correlation between indicators (pTM, pLDDT, and ipAE) and RMSD_Cα metrics. (G-I) Relationship between residue length and RMSD metrics.

#### The RMSD-based evaluation of non-canonical cyclic peptide complexes

The RMSD-based evaluation of AlphaFold3-predicted cyclic peptide complexes was similar to that of monomer structures. In addition to the cyclic peptide ligand structures, we investigated AlphaFold3’s performance in predicting the target protein structure. As shown in Figure 4B, the RMSD_Cα values of the target protein ranged from 0.241 to 1.130 Å, with an average RMSD value of 0.549 Å. These targets contain different conformations (Figure 4D), demonstrating the model’s generality. But considering the overly low RMSD values for these long chains, we reasoned that it may be attributed to the fact that the training dataset of AlphaFold3 contains these targets, therefore leading to accurately predicting the corresponding conformation.

For the ligand, the average RMSD_Cα, RMSD_aa, and RMSD_uaa reached 0.592, 1.105 and 1.121 Å (Figure 4C). Take the 5IV2 as an example, the RMSD_Cα, RMSD_aa, and RMSD_uaa of the predicted ligand of this complex were 0.372, 0.466 and 0.353 Å, respectively, which were closely similar to the native fold. Notably, we observed reduced RMSD values for ligand-bound conformations compared to those of the monomers. This phenomenon likely resulted from structural constraints imposed by the target protein’s binding interface, which restricted the conformational sampling of the cyclic peptide. Namely, AlphaFold3 demonstrates superior predictive performance for cyclic peptide complexes than monomers.

#### The relation between ligand_RMSD and indicators

In cyclic peptide complex prediction, ipAE and ipTM serve as commonly used assessment metrics for evaluating structural plausibility, with widespread adoption in peptide design for structure validation. Figures 4E-G demonstrate the relationship between these metrics and ligand RMSD values. While no statistically significant correlation was observed in the analysis, the predictive reliability increased when ipAE values fell below 0.1 (Figure 4G). Structures satisfying ipAE<0.1 exhibited significantly lower Cα-RMSD values (below 0.5 Å) compared to those above this threshold. However, the ipAE < 0.1 criterion represents a relatively stringent metric. It should be noted that the constrained cyclic architecture inherently limits the conformational flexibility of these molecules. Consequently, establishing a reliable ipAE threshold for cyclic peptides remains an ongoing challenge that will require further investigation and experimental validation.

As for the ligand_pLDDT, the conclusion about pLDDT is consistent with that in the experiment focusing on cyclic peptide monomers. Our analysis reveals a positive correlation between pLDDT values and predicted structure plausibility, with higher confidence scores corresponding to more reliable structural predictions. However, as demonstrated in Figures 4E and 4H, our benchmarking data do not support the use of pTM as a reliable metric for assessing prediction accuracy in cyclic peptide complexes.

In summary, the ipAE and pLDDT metrics can serve as preliminary structural assessment tools, given the current lack of standardized evaluation criteria. However, comprehensive fold validation still requires integrative expert analysis that combines biochemical principles with structural bioinformatics approaches. Regarding ipTM, this metric does not reliably reflect the accuracy of predicted cyclic peptide structures.

### The Effect of MSA and Template Information in the Alphafold3 Model

One of the primary modifications in AlphaFold3 compared to AlphaFold2 is the change on MSA processing, achieved through a smaller and simpler MSA embedding block. This adjustment aims to accommodate a diverse range of chemical entities without excessive reliance on special cases. In previous work, an ablation experiment demonstrated that MSA information played a critical role in modeling protein conformation. In this study, we also investigated whether MSA features can enhance AlphaFold3’s ability to predict cyclic peptide structures. To address this question, we removed the MSA feature input from this model and evaluated AlphaFold3’s performance in predicting cyclic peptide monomer structures. As shown in Figure 5, the AlphaFold3_no_MSA RMSD values of Cα, all atoms, and unnatural atoms (1.856, 3.178 and 2.908 Å) were not significantly different from the original AlphaFold3. In other words, the impact of MSA information on the accuracy of predicted structures is relatively limited.

**Figure 5.**
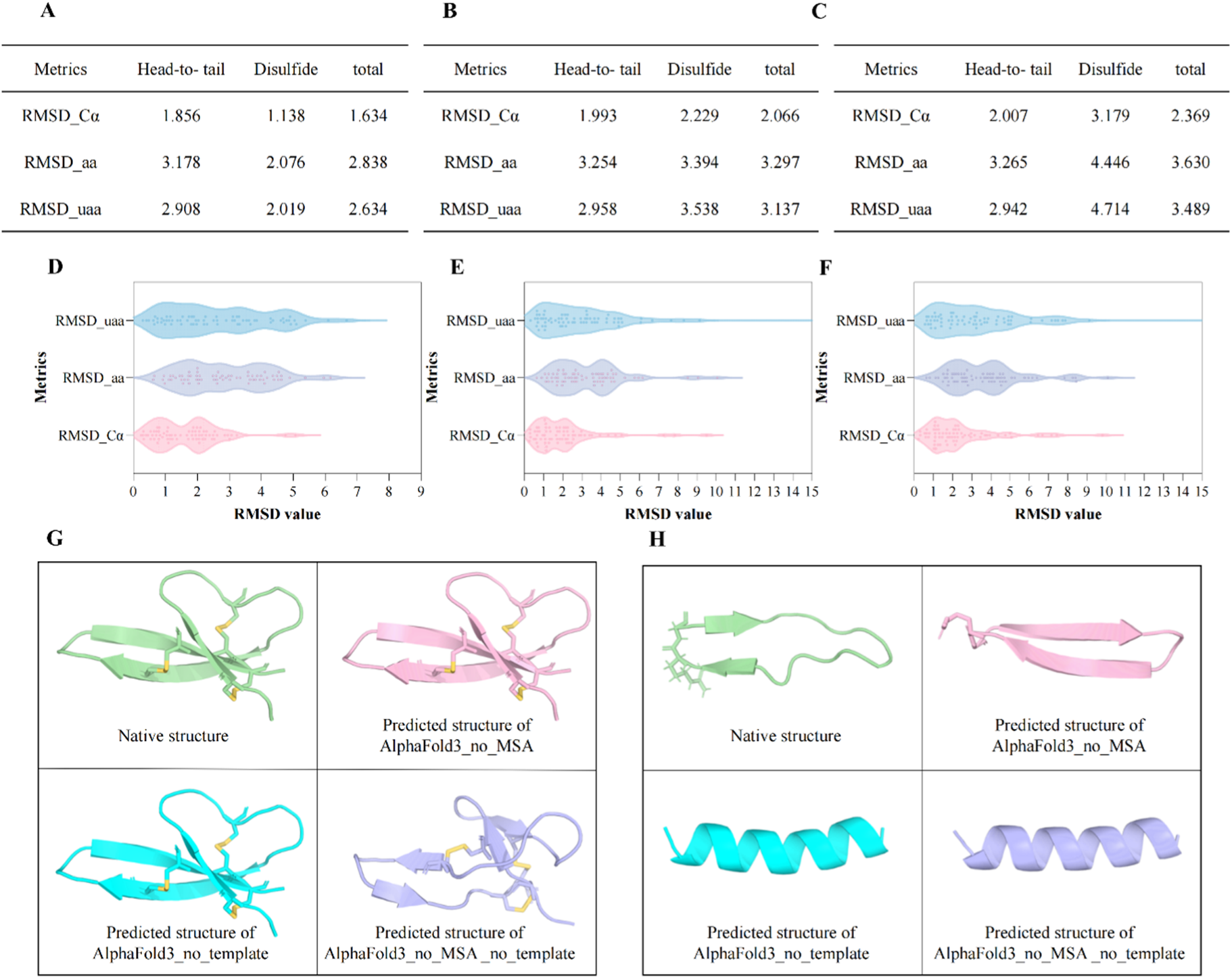
The ablation experiment of AlphaFold3 in predicting non-canonical cyclic peptide monomers. (A) The average RMSD values of cyclic peptide monomers predicted by AlphaFold3_no_MSA. (B) The average RMSD values of cyclic peptide monomers predicted by AlphaFold3_no_template. (C) The average RMSD values of cyclic peptide monomers predicted by AlphaFold3_no_MSA_no_template. (D) The distribution of RMSD values of cyclic peptide monomers predicted by AlphaFold3_no_MSA. (E) The distribution of RMSD values of cyclic peptide monomers predicted by AlphaFold3_no_template. (F) The distribution of RMSD values of cyclic peptide monomers predicted by AlphaFold3_no_MSA_no_template. (G) The predicted structures of 4E82 predicted by AlphaFold3 variants compared to the native structure. (H) The predicted structures of 2M2X predicted by AlphaFold3 variants compared to the native structure.

In contrast, template feature removal resulted in more substantial performance degradation. As shown in Figure 5B, the exclusion of template information increased the average RMSD_Cα to 2.066 Å. Particularly pronounced accuracy reductions were observed in the disulfide cyclic peptide predictions (RMSD_Cα values increase from 1.203 to 2.229 Å). As for the AlphaFold3_no_MSA_no_template prediction, such performance decrease is more evident, the RMSD_Cα, RMSD_aa and RMSD_uaa dropped by 0.698, 0.768 and 0.897 Å, respectively. Additionally, the absence of template and MSA resulted in a great performance decrease of AlphaFold3 in predicting disulfide cyclic peptides. The RMSD_Cα, RMSD_aa and RMSD_uaa dropped to 3.179, 4.446 and 4.714, meaning complete failure in this work.

These findings establish template information as a critical factor of AlphaFold3’s predictive capability for non-canonical cyclic peptides. For practical applications under computational resource constraints or requiring accelerated prediction speeds, we recommend retaining template information while optionally removing MSA features.

### Mechanistic Insights into Alphafold3’s Cyclic Peptide Prediction

Notwithstanding the explicit input of cyclization constraints, we observed structural violations in a subset of predictions where the backbone failed to form closed cyclic conformations. As displayed in Figure 6A, these non-cyclized structures exhibited discontinuous connectivity between terminal residues despite the defined cyclization information. To better understand this limitation, we sought to examine the underlying mechanisms governing AlphaFold3’s cyclic peptide predictions.

**Figure 6.**
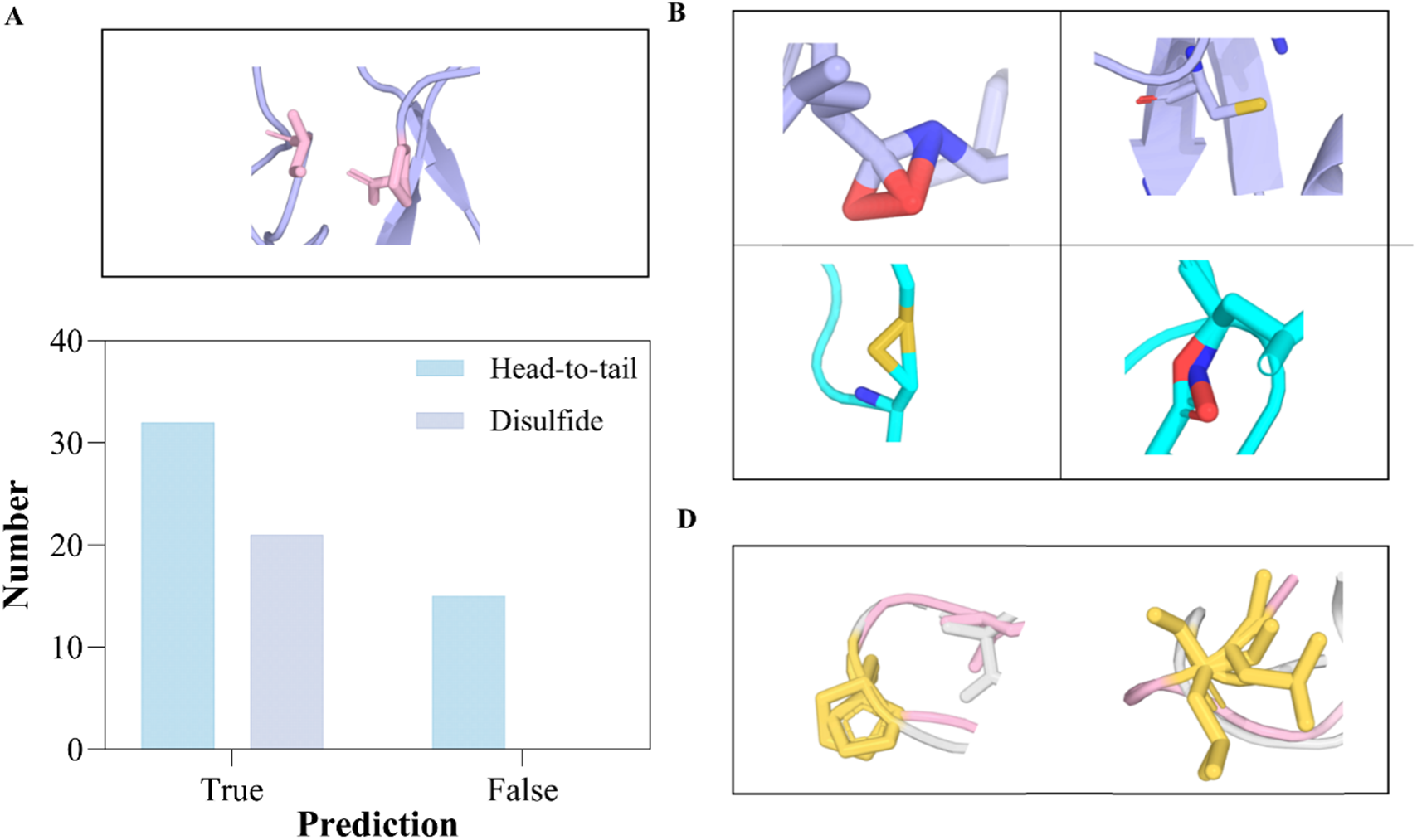
The mistakes of AlpahFold3 in predicting non-canonical cyclic peptides (A) non-cyclized structures (B) different atom-level structure mistakes. (C) charity errors.

Previous implementations enabling AlphaFold to predict cyclic peptide structures relied on incorporating conformational constraints into its positional encoding matrix. This approach successfully guided the AlphaFold model to generate cyclized folds. However, our investigation revealed that AlphaFold3 does not employ this methodology for cyclic peptide prediction. Furthermore, our examination of the source code indicates this information is not explicitly integrated into the modeling process. Consequently, the probability of achieving a cyclic conformation is not strongly influenced by user-defined constraints. Instead, the resulting conformation (whether linear or cyclic) appears predominantly determined by the model’s learned patterns from training data rather than explicit input constraints. This suggests AlphaFold3 lacks a deterministic mechanism to enforce cyclic conformations during structure prediction.

To validate our observations, we systematically analyzed AlphaFold3’s predictions for the monomer subset and identified such failures in this experiment. While these structures closely resembled native conformations, the peptide termini failed to connect and form cyclic structures, instead maintaining linear configurations. Among 57 head-to-tail cyclic peptide monomers, 15 were incorrectly predicted as linear. For disulfide-bridged cyclic monomers, all predictions showed the cyclization.

This discrepancy in prediction accuracy of head-to-tail and disulfide cyclic peptides is attributed to structural similarities between disulfide cyclic peptides and conventional linear peptides, both requiring disulfide bonds for stability. In essence, disulfide cyclic peptides can be regarded as a specialized subclass of linear peptides, explaining AlphaFold3’s relatively better performance on these structures compared to head-to-tail cyclic peptides (Figure 6A). Furthermore, we identified specific structural mistakes in the predicted cyclic conformations, such as atomic clashes in head-to-tail cyclic peptides, and incorrect disulfide bond geometries between cysteine residues. Representative examples of these errors are presented in Figure 6B. These findings further demonstrate that AlphaFold3 still exhibits limitations in accurately modeling atomic-level structural details, while showing relatively stronger performance in predicting global cyclic peptide conformations. This observed phenomenon may be associated with AlphaFold3’s removal of traditional relax module, which could introduce atomic-level structural clashes. However, further investigation is required to establish a definitive relationship between the model architectural change and the observed structural problems.

Recent studies have demonstrated AlphaFold3’s limitations in accurately predicting atomic chirality^33^. In this work, we further investigated AlphaFold3’s predictive accuracy regarding chiral centers. No surprise, the AlphaFold3 can’t distinguish the chirality of residues and always predict the D-amino residues to be L-amino residues. Two examples are displayed in Figure 6C.

To avoid such biases or inaccuracies, users should carefully check the model’s predicted structures before conducting further analyses or experiments. Furthermore, an effective input should be added to the AlphaFold3 model, allowing information about cyclization to be incorporated, thereby generating conformations that meet the input requirements. For researchers requiring large-scale cyclic peptide structures, we recommend employing an automated preprocessing pipeline to enhance reliability.

## Conclusion

While AlphaFold3 has demonstrated significant advancements in protein structure prediction involving unnatural residues and small molecules, its applicability to non-canonical cyclic peptide structural determination remains inadequately validated. In this systematic investigation, we performed comprehensive benchmarking using a structurally diverse non-canonical cyclic peptide dataset, including monomers and complexes to evaluate the model’s efficacy in this specialized domain. Our investigation also revealed an accessibility barrier for researchers attempting large-scale cyclic peptide prediction, prompting us to develop an automated computational pipeline that accelerate the prediction workflow.

Quantitative analysis using three RMSD metrics revealed the model’s fundamental applicability for cyclic peptides with average RMSD_Cα (1.671 Å), RMSD_aa (2.862 Å), and RMSD_uaa (2.592 Å). Based on these systematic evaluations, we provide the following critical implementation guidelines:

1. Confidence metric reliability: Although pLDDT does not exhibit a clear linear correlation with prediction accuracy, higher pLDDT values generally correspond to an increased likelihood of correctly predicted non-canonical cyclic peptide structures. Therefore, pLDDT can serve as a preliminary assessment metric for evaluating structural predictions generated by AlphaFold3. Additionally, ipAE values also provide a reliable scoring measure for assessing predicted cyclic peptide complexes. In contrast, our benchmarking experiments demonstrate that ipTM is not a robust metric for evaluating cyclic peptide prediction accuracy.
2. The importance of template and MSA: Controlled ablation experiments demonstrate the impact of template structures and MSAs. We recommend maximizing the incorporation of available biological context through these inputs when computationally feasible. If resources are limited, we recommend maintaining the template information rather than the MSA information.
3. The AlphaFold bias: Compared to head-to-tail cyclic peptides, AlphaFold3 demonstrates superior predictive accuracy for disulfide cyclic peptides. Furthermore, AlphaFold3 exhibits stronger modeling capability for backbone atoms than for side chain or unnatural residue components. Finally, we observed that cyclic connectivity constraints are not explicitly encoded in AlphaFold’s feature embedding process, necessitating careful validation of predicted structures.

This study conducts the first comprehensive benchmarking experiment for applying AlphaFold3 to non-canonical cyclic peptide modeling with practical implementation guidelines. Our findings and recommendations aim to accelerate research in cyclic peptide structural biology by enabling more effective utilization of this powerful prediction tool.

## Acknowledgment

This project was supported by the Macao Science and Technology Development Fund (Grant No. 0151/2024/RIA2) and the internal grant from Macao Polytechnic University (RP/FCA-07/2024).

